# Optogenetic Control of the Integrated Stress Response Limits Glioblastoma Invasion

**DOI:** 10.1101/2025.08.22.671866

**Authors:** Lisa K. Månsson, Ethan Dickson, Lun Hao, Angela A. Pitenis, Maxwell Z. Wilson

## Abstract

The integrated stress response (ISR) is a highly conserved cellular signaling network, allowing cells to adapt and respond to various stressors. In one of the toughest cancers to date, glioblastoma multiforme (GBM) with aggressive spread and high recurrence rates, the role of the ISR is yet to be well understood - whether activation may suppress or promote this disease – and drug-treatment of GBM has thus far shown inconclusive results. In this work we use an optogenetic tool, opto-PKR, to specifically trigger ISR-activation with light with high spatiotemporal control via the PKR-kinase, avoiding potential upstream damage or side effects from drugs. Using immunofluorescence and RNA-sequencing we show that targeted ISR-activation reaches levels where both adaptive (ATF4) and terminal response (CHOP) of the ISR are activated, which show downregulation of genes associated with extracellular environment and glial cell migration, further supported by ECM-stain and scratch assays. Further, we show inhibition of aggressive spread for ISR-activated GBM spheroids in collagen 3D culture. Photopatterning of ISR-activation in spheroids demonstrates a cell intrinsic effect at tissue scale, and recovery studies indicate a tunable, non-ablative intervention space. These findings suggest a route to containment and motivate ISR-activating small molecule screening in GBM models.

## 1. Introduction

When a cell is exposed to an environmental or pathological condition, such as cancer, the Integrated Stress Response (ISR) allows the cell to adapt and respond through transcriptional changes [1]. The ISR is activated by four kinases (PERK, PKR, HRI, and GCN2), each of which will become active when clustered with multiples of itself. Clustering of the different kinases is induced by different cellular stressors, such as nutrient deficiency, oxidative stress, double stranded RNA present in viral infection, or misfolded proteins. Once activated, the kinases will start phosphorylating the α-subunit a transcription factor called eukaryotic translation initiation factor (eIF2α). In its phosphorylated state eIF2 is unable to be part of a ternary complex that is a crucial component in protein translation, thus shutting down majority of translation in the cell. Simultaneously, phosphorylated eIF2 promotes selective protein translation, including downstream ISR transcription factors; ATF4, associated with an adaptive stress response, and CHOP, activating terminal stress response towards cell death [2], [3], [4]. The ISR also activates negative feedback loops, translating the transcription factor GADD34 [5], [6] that together with protein phosphatase 1 (PP1) and globular actin (G-actin) forms a complex that de-phosphorylates eIF2 [7], [8]. In this way, activation of the downstream ISR transcription factors helps the cell respond and adapt to stressful conditions, and in severe cases undergo apoptosis.

As the ISR has this big impact on the programming of the cell, it is to no surprise it is an essential signaling network for mammalian development and survival [9], [10], [11]. If not lethal, milder dysregulation of the ISR has been correlated with multiple different diseases, including metabolic disorders [9], [12], [13], [14], [15], multiple intellectual disabilities [16], [17], [18], [19], [20], [21], [22], cognitive and neurodegenerative disorders [23], [24], [25], [26], as well as long-term memory formation [1], [27], [28], [29], [30], [31], [32], [33], [34], which is highly dependent on protein synthesis [35], [36], [37], [38], [39]. While multiple studies connecting ISR to disease show that ISR-inhibition ameliorates pathogenesis [40], [41], [42], [43], [44]. Others show ameliorated states from ISR-activation [45], [46], [47], [48], [49], [50], [51]. These results show the complexity of the ISR signaling network and how it may act differently in different cells and diseases.

The ISR also plays a big role in different cancers, as tumor cells are using it to overcome oncogenic and therapeutic stressors [52], [53], [54]. Increased protein synthesis, metabolic pressure [55], [56], genomic instability as well as decreased nutrient and oxygen availability [57] are all different oncogenic stressors for the cell to cope with during tumor progression, why multiple cancers (e.g., prostate cancer [58], [59], [60], lung adenocarcinoma [61], lymphomas [62], melanoma and pancreatic tumors [63]) have been found to use the ISR for survival, as a main regulator of adaptation to stress. In addition, ISR-activation has also been found shielding cancer cells from treatment [64], [65] or drive apoptosis of cancer cells. Examples include drugs driving ISR-activation from nutritional stress [66], [67], [68], drugs accelerating misfolded protein degradation thus activating the ISR [69], [70], or through amplified activation via inhibition of negative feedback of the ISR in different cancers [71], [72], [73].

With less than 7% five-year post-surgery survival rate [74], Glioblastoma Multiforme (GBM) is one of the most challenging cancers of today because of its aggressive growth and shielding mechanisms that make treatments less efficient [1]. GBM tumors are typically located in the frontal lobe, often found in the sensitive broca area, the region for language and cognition, and spread with finger-like protrusions and single cancer cells migrating long distances from their origin (**Fig. 1a**) [75]. As GBM progresses, the extracellular environment also changes and is getting denser, acting as a shield around the tumor that makes common cancer treatments inefficient [76]. When surgery becomes the main option of treatment, the aggressive spread makes it hard to safely and fully remove the tumor, and thus the common return of the disease [77].

**Figure 1.**
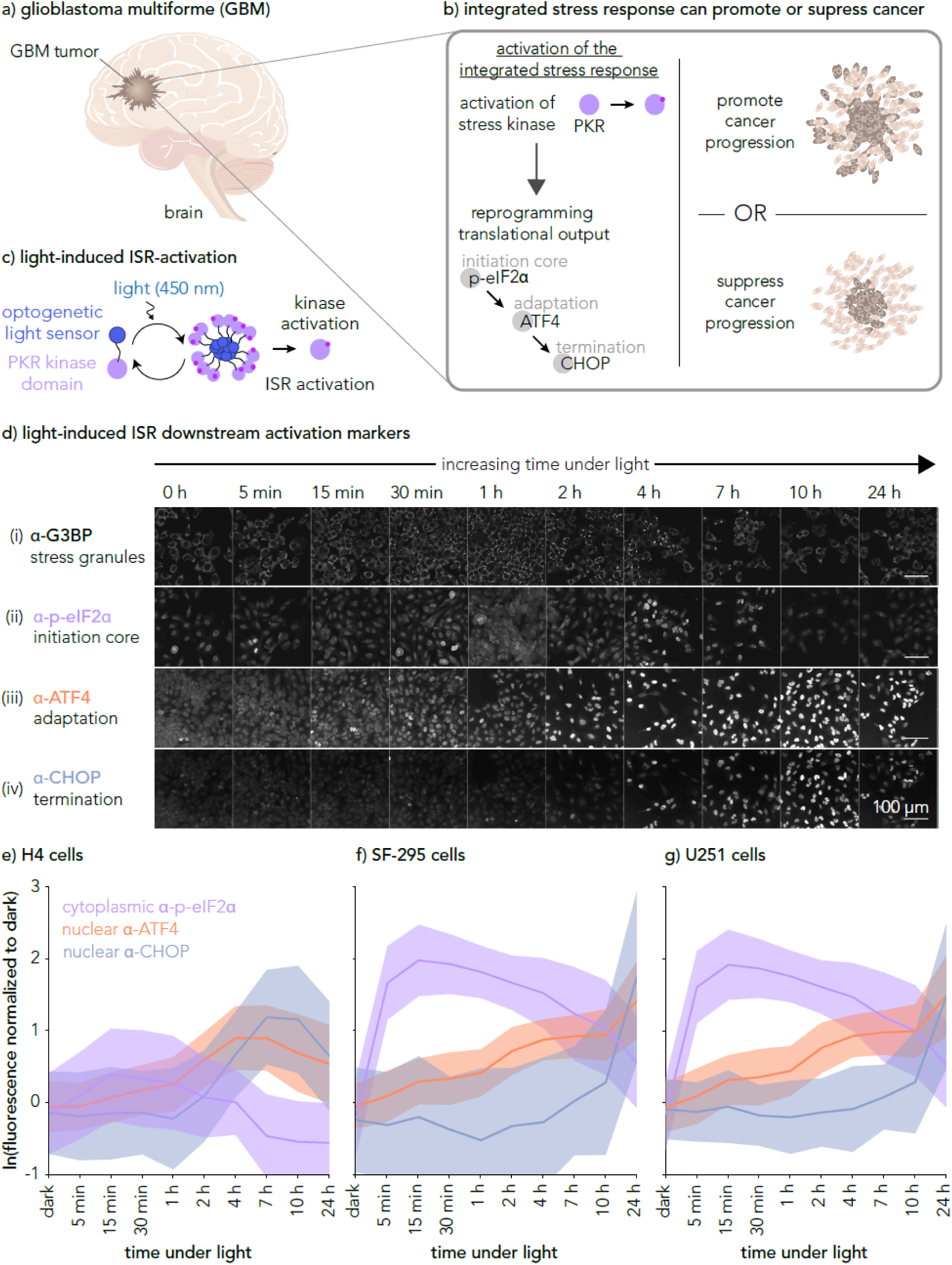
Opto-PKR allows for high spatiotemporal control with light-triggered ISR-activation to examine its role in GBM cancer progression. a) GBM is a severe type of brain cancer with aggressive spread; finger-like protrusions and single cancer cells migrating long distances from their origin. b) ISR is a complex signaling network that reprograms translational output in the cell via phosphorylation of eIF2α by the PKR kinase, followed by activation of transcription factors ATF4 and CHOP, through which the cell adapts and responds to the stress applied. ISR-activation has been suggested to promote but also to suppress GBM progression in different studies. c) Using an optogenetic tool for ISR-activation developed by Batjargal et al. [90], we achieve reduced molecular complexity and high spatiotemporal control of PKR-kinase activation inducing artificial ISR-activation in cells. d) Immunofluorescence images of downstream ISR-markers in H4 neuroglioma cells after different amount of ISR-activation (time under light). e), f), and g) Quantification plots based on immunofluorescence images of cytoplasmic α-p-eIF2α, nuclear α-ATF4, and nuclear α-CHOP, for three different cell lines; H4 (e), SF-295 (f), and U251 (g). All fluorescence values are normalized to the dark condition average for each protein. More than 100 cells were quantified for each time point for each cell line to obtain means (solid line) and standard deviation (shaded regions). Representative immunofluorescence images for SF-295 and U251 are shown in **Supplementary Fig. 1** and **2**.

The role of the ISR in GBM is yet to be well understood - whether activation may suppress or promote this disease. Studies treating GBM cells with different drugs activating parts of the ISR signaling network have thus far shown inconclusive results; both pro- and anti-cancer progression effects (**Fig. 1b**) [78], [79], [80], [81], [82], [83], [84]. ATF4 is not only part of the adaptive response of the ISR (**Fig. 1b**) but also in many other signaling networks (e.g., WNT, mTOR [85]) and has in recent work been seen to support cancer progression by helping GBM cells adapt to the stressful tumor environment (e.g., activation versus knockout studies of ATF4 [78], [82], [86], and the ATF4-SPHK1 cascade [79]). However, in other studies examining the effect of different drugs on GBM it was shown that the drug which suppressed cancer progression *in vivo* (e.g., Nootkatone [84], and Withaferin A [83]) activated the ATF4-CHOP signaling cascade. In addition, ISR-activation in combination with one of the most common GBM drugs temozolomide has been seen to enhance its efficiency [87], [88], [89]. Common for the studies showing suppression of GBM with ISR-activation, is that the terminal transcription factor CHOP was always upregulated as part of the activation. Considering these findings, we hypothesized that the fate of suppression or promotion of GBM may be dependent on the level of ISR-activation. Specifically, we aimed to examine if high levels of ISR-activation, to the point where both ATF4 and CHOP are activated, may be a target for GBM suppression.

An optogenetic tool for ISR-activation allows one to study the process dynamics and feedback controls of this complex signaling network in isolation from confounding upstream damage or drugs (**Fig. 1c-g**). In contrast to a real stressor activation, the light input is a tunable activator allowing complex design of input dynamics, duration and intensity, as well as high spatiotemporal control of the ISR-activation.

## 2. Materials and Methods

### 2.1. Cell Lines

The cell lines used in this work include H4, derived from neuroglioma, U251, and SF-295. The latter two are derived from GBM and part of the NCI-60 human tumor cell line screen for cancer research [91].

For H4 cells, wild type (WT) as well as the genetically modified H4 opto-PKR cell line with light-activated ISR were used [90]. Cells were handled according to standard protocols, incubated in 37°C with 5% CO_2_, passed every 3-4 days, and grown in DMEM supplemented with 10% Fetal Bovine Serum and 1% penicillin-streptomycin.

U251 cells were grown in DMEM media while SF-295 were grown in RPMI 1640 media, both of which were supplemented with 10% Fetal Bovine Serum and 1% penicillin-streptomycin. Both cell lines were handled according to standard protocols, incubated in 37 °C with 5% CO_2_, and passed every 3-4 days. U251 and SF-295 cell lines were transduced to express the same opto-PKR protein as the H4. Both were selected with 2 ug/mL of Puromycin for 5 days.

All opto-PKR cell lines were always handled in the dark or red light to not ISR-activation outside stimulation experiments.

### 2.2. Immunofluorescence of ISR-activation Markers

For all three cell lines, both WT and opto-PKR cells were plated around 15 000 cells per well in fibronectin-coated glass 96-well plate a day before start of stimulation experiment. Using the Optoplate [92] with a thin diffuser on top, cells were stimulated in the well plate over the course of 24 hours, with pulsed light sequences of 5 s ON, 15 s OFF in 450 nm blue light of 200 μW/cm^2^. Stimulation conditions were defined by varying the time of light exposure (0 min, 5 min, 15 min, 30 min, 1 h, 2 h, 4 h, 7 h, 10 h, and 24 h) during the 24 h experiment realized by a programmed delay in the start of light pulses for different wells. After stimulation, cells were fixed in 4% paraformaldehyde solution for 20 minutes, thereafter, washed with DPBS before staining. Cells were first stained with primary antibodies in blocking buffer (0.05 w/v % saponin, 0.5 w/v % BSA, 50 mM ammonium chloride, and 0.02 v/v % NaN_3_, in 1× PBS, pH-adjusted to pH 7.2-7.4, and filtered through 0.22 μm sterile PVDF filter before stored at 4 °C) overnight for 20-28 hours in 4°C. Half of the wells were incubated with α-p-eIF2α (Abcam, cat no. ab32157, rabbit), and α-G3BP (BD Biosciences, cat no. 611127, mouse), while the other half got α-ATF4 (Cell Signaling Technology, cat no. 11815S, rabbit), and α-CHOP (Cell Signaling Technology, cat no. 2895, mouse). Thereafter cells were washed in T-PBS (0.1 v/v % Tween 20 (Fisher Scientific, cat no. BP337-500) in 1× PBS) three times (first instant, second with 20 minutes incubation, third instant) before adding blocking buffer with secondary antibodies and DAPI for nuclear stain, incubated at RT for 30 min to 1 h with the well plate covered for reduced light exposure. After secondary incubation cells were once again washed three times with T-PBS, as previously described, and finally left in 1× PBS. With stained cells in PBS, all wells were imaged with the Voyager CQ1 confocal imager (Yokogawa), saving maximum intensity projection images of a 10 μm z-stack, with a z-step of 1 μm, around the best focus of the nuclear stain (DAPI).

For quantification, images were analyzed in Python using CellPose [93], [94] to segment each cell outline as well as its nucleus. The nuclear fluorescence intensity was measured for α-ATF4 and α-CHOP to study their activation over time. For phosphorylation of eIF2α the cytoplasmic fluorescence intensity was computed by subtracting the segmented area of the nuclei from the whole cell mask to create a cytoplasmic mask without nuclear fluorescence.

For H4 cells this experiment was repeated three times, and for U251 and SF-295 the experiment was performed once. For each repeat, each time point for each cell line include quantification of a minimum of 100 individual cells.

### 2.3. RNA-sequencing and Analysis

H4 opto-PKR cells were plated around 15 000 cells per well in a fibronectin-coated glass 96-wells the day before start of stimulation experiment. Using the Optoplate [92] with a thin diffuser on top, cells were stimulated in the well plate over the course of 24 hours, with pulsed light sequences of 5 s ON, 15 s OFF in 450 nm blue light of 200 μW/cm^2^. Control wells were kept in the dark. After stimulation, cells were collected using extraction buffer from RNeasy Mini Kit (Qiagen, cat no. 74104) and kept at -80 °C until RNA-extraction was performed with the same kit following sample preparation instructions. Concentration of the RNA samples were measured with a nanodrop, thereafter diluted to 5 ng/μL before library preparation and sequencing following previously developed protocols for bulk cell sequencing [90], adapted from the single-cell sequencing method “CEL-Seq2” [95], [96].

RNA-sequencing data from all samples were grouped by condition and the 24 h light stimulation (N=3 samples from individual wells) data was compared to that of 24 h dark (N=2 samples from individual wells). Differential gene expression analysis was performed in R (version 4.4.0) using the package *DEseq2* (version 1.44.0) [95], [97]. A total of 17,074 genes were detected, thereafter, filtered on low counts (less than 10 counts as total within the condition or genes with less than 10 counts in a single sample), reducing the set to 15,035 genes. Applying differential gene expression analysis with *DEseq* using a significance level of α = 0.05, a total of 1,905 genes came out as differentially expressed, where just over half of the genes (987) were upregulated and the other half (918) were downregulated. To avoid false positives in the continued analysis, log fold change values of genes with low counts were shrunk using *lfcShrink* (type: ashr, α = 0.05) [97]. The difference in fold change for the gene set before and after shrinkage is visualized in MA-plots in **Supplementary** Fig. 3. Correlation between replicate samples as well as between categories were analyzed using Pearson correlation, presented in **Supplementary** Fig. 4. Finally, gene enrichment analysis was performed on the 25 most up and downregulated genes using g:profiler [98].

### 2.4. ECM-stain Assay

Both WT and opto-PKR H4 cells were seeded around 60 000 cells per well in fibronectin-coated glass bottom 24-well plates the day before the start of the stimulation experiment. Cell culture media was exchanged in all wells just before the start of the light stimulation. For ISRIB control conditions, media was supplemented with ISRIB (MedChemExpress) [99], [100] at a working concentration of 200 nM.

Using the Optoplate [92] with a thin diffuser on top, cells were stimulated in the well plate over the course of 72 hours, with pulsed light sequences of 5 s ON, 15 s OFF in 450 nm blue light of 200 μW/cm^2^. After stimulation, cells were fixed in 4% paraformaldehyde solution for 20 minutes, then washed with DPBS before staining. Sirius Red/Fast Green Collagen Staining Kit (Chondrex #9046) was used to stain fibrillar collagens and non-collagenous ECM, following the instructions of the kit. With stained cells in PBS, all wells were fully imaged at 4× magnification in a 4×4 grid of positions with Voyager CQ1 confocal imager (Yokogawa), saving z-sum images of a 280 μm z-stack, with a z-step of 4 μm between each image. This range was set based on ECM dye fluorescence in z, capturing the full fluorescence-intense range. To obtain quantitative data extraction buffer from the same kit was used to extract all ECM-bound dye in a well and moved into a sterile plastic 24-well plate for absorption reading in Cytation 5 imaging reader (BioTek) from which mass estimates of each component (Sirius Red and Fast Green) were calculated, following instructions from the kit. The fraction of Sirius Red to Fast Green per well was calculated to obtain the collagen fraction compared to non-collagenous ECM. The full experiment was repeated three times.

After ECM stain assay was performed, the cells were stained with DAPI and imaged in the same way as described for the ECM stain kit above to obtain images for quantification of nuclear area coverage per well.

### 2.5. Scratch Assay

Both WT and opto-PKR H4 cells were seeded around 500 000 cells per well in plastic 12-well plates the day before start of stimulation experiment. A scratch was made in each well using the back side of a sterile 200 μL pipette tip. All wells were thereafter rinsed with DPBS, before cell culture media was added just before the start of the light-stimulation. For ISRIB control conditions, media was supplemented with ISRIB at a working concentration of 200 nM. Using a custom-built light box with blue LEDs at the bottom and multiple diffusers layered on top, cells were stimulated in the well plate over the course of 72 hours, with pulsed light sequences of 5 s ON, 15 s OFF of about 200 μW/cm^2^ light intensity. A duplicate well plate was wrapped in aluminum foil and kept in dark throughout the full experiment as control. The full scratch in each well was imaged at 4× magnification at time point 0 h (just before stimulation start), and then repeatedly after 24 h, 48 h, and 72 h. Image analysis was performed in ImageJ Fiji [101], [102], where each scratch width was measured in at least ten different positions to obtain a mean value per time point per condition. The full experiment was repeated three times.

### 2.6. 3D Spheroid Culture

Spheroids were generated from both opto-PKR and WT cells of the H4, U251, and SF-295 cell lines, by seeding 5,000 cells per well in low-attachment 96-well plates (Thermo Scientific™ Nunclon™ Sphera™) and left in the incubator for about 36 hours. For the last 12 hours live cell stain CellTracker Red CMTPX (CellTracker™ Fluorescent Probes) was added to the spheroids at a final concentration of 5 μM in cell culture media via partial media exchange in the low-attachment wells. Thereafter, cells were gently washed with DPBS through three partial media exchanges to remove any dye residue, left to rest in the incubator. Rat Tail Collagen I (3 mg/mL, Corning) acidic stock was neutralized with 2.5 v/v % 1M NaOH and diluted to a concentration of 2 mg/mL in cell culture media to pre-coat a glass bottom 96-well plate kept on ice (to avoid gelation) with 40 μL neutralized collagen matrix (2 mg/mL) per well. To evenly distribute the gel in each well and remove air bubbles the well plate was centrifuged at 970×g for 2 minutes at 4 °C. The pre-coated 96-well plate was then kept in RT while another 450 μL of neutralized collagen (2 mg/mL) was prepared as stock. For each spheroid, 40 μL of the collagen stock was aliquoted into a test tube to which the spheroid was transferred from the low-attachment well plate using careful pipetting. The 40 μL of collagen with the spheroid was thereafter transferred into one of the pre-coated wells in the glass bottom 96-well plate. Once all spheroids were plated in collagen, the well plate was put in the 37 °C incubator for one hour to allow gelation of the collagen, before cell culture media was added on top. From here, spheroids were either studied under time lapse imaging for 24 h or imaged at discrete time points before and after 24 h of stimulation.

For time lapse imaging, the well plate with spheroids was put for 24 h on a SoRa spinning disc confocal microscope (Nikon), with environmental control of maintaining humidity, 37 °C, and 5% CO_2_. Using the software plugin *Jobs*, a number of wells were selected for stimulation with blue light using the 488 nm laser, exposing spheroids to 500 ms of light every 2 minutes with a laser power of 20%, focused on the middle z-plane of the spheroid. All spheroids were imaged taking a z-stack of 600 μm (z-step 20 μm) in the 561 nm channel (600 ms exposure time, 100% laser power), visible with the CellTracker Red stain, at every hour of the time lapse.

For discrete time point imaging, all spheroids were imaged before the start of the light stimulation taking a z-stack of 600 μm (z-step 20 μm) in the 561 nm channel (600 ms exposure time, 100% laser power), visible with the CellTracker Red stain, as well as a middle z-plane imaging in DIC. Thereafter, the plate was put on the Optoplate, programmed to stimulate selected wells with blue (450 nm) LED of 200-500 μW/cm^2^ at a repeated pulsed illumination of 5 s ON, 15 s OFF. After 24 h spheroids were once again imaged as described above. To study recovery after ISR-activation, spheroids were kept in the dark for 48 h and imaged again.

Manual quantification of spheroid morphology was performed in Fiji [101], tracing the spheroid perimeter, thereby obtaining measurements of perimeter, area, and circularity. Spheroid circularity, *c*, was calculated as defined by Eq. (1),

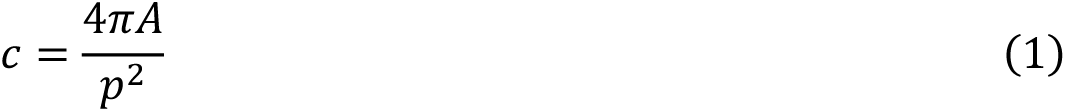

where *A* is the spheroid area, and *p* is the spheroid perimeter. For a perfectly circular object *c*=1, meanwhile protrusions from spheroids drastically decrease *c* to a value less than one. For time lapse quantifications, each spheroid at each time point was circled n=3 times to obtain average values. For endpoint experiment quantifications, each spheroid was circled once.

### 2.7. Photopatterned 3D Spheroid Culture

Spheroids were generated from both opto-PKR and WT cells of the SF-295 cell lines as described above. The well plate with spheroids was put for 22 h on a SoRa spinning disc confocal microscope (Nikon), with environmental control of maintaining humidity, 37 °C, and 5% CO_2_. Using digital micromirror devices (Mightex Polygon 400), a rectangular region of interest was chosen for blue light illumination covering the right half of a spheroid in the middle z-plane of the 3D-structure. In the software Jobs, wells were selected for stimulation with blue light using the blue light (447 nm) X-Cite xLED (Excelitas Technologies) connected to the DMD, exposing spheroids to 500 ms of light every 2 minutes with LED power at 50%, focused on the middle z-plane of the spheroid. All spheroids were imaged taking a z-stack of 600 μm (z-step 20 μm) on the 561 nm channel (600 ms exposure time, 100% laser power), visible with the CellTracker Red stain, at every hour of the time lapse.

## 3. Results

The ISR is activated by four kinases (PERK, PKR, HRI, and GCN2), each of which will become active when clustered with multiples of itself. With optogenetic clustering tools, we can engineer cells who induce this activation in response to light. Batjargal *et al.* developed an optogenetic tool for light-induced ISR activation via Protein Kinase R (PKR) -clustering, “opto-PKR” [90]. By fusing PKR to the cry2olig optogenetic tool, a blue light oligomerizer [103], cells are endowed with light-inducible downstream ISR-activation, greatly reducing the combinatorial complexity of signals turned on by real stressors. In this work, the same optogenetic tool was used to study the relation between ISR-activation and disease.

### 3.1. Opto-PKR Enable Light-Induced ISR-activation

Using engineered light-stimulation devices [92], we achieve high spatiotemporal control over cellular stress in addition to intensity and pulse modulation and can spatiotemporally control ISR activation in cells. **Fig. 1d-g** and **Supplementary** Fig. 1 and 2 show the 24 h time-resolved downstream ISR activation for cell lines H4, SF-295, and U251 under stress using light-activation of the opto-tool: stress granule formation (G3BP), the ISR initiation core in phosphorylation of eIF2α (p-eIF2α), followed by adaptive transcription factor 4 (ATF4), and finally terminal ISR-response represented by transcription factor CHOP. The observed protein dynamics are quantified and indicate clear cellular stress under light-induced ISR-activation, in reference to previous literature, including stress response to drugs [90]. Wild type cells (not shown) under the same illumination conditions did not show ISR-activation in either of the four stress markers. These results support that light-activated PKR-clustering with opto-PKR triggers spatiotemporally controlled ISR-activation in all three cell lines chosen.

### 3.2. The Integrated Stress Response Changes Gene Expression in Cell Survival and Death

To examine if the ISR may be a target signaling network for brain cancer regulation, RNA sequencing was performed in opto-PKR H4 cells exposed to 24 h of light (N=3), compared to control samples in the dark for the same amount of time (N=2) (**Fig. 2a**), showing clear changes in gene expression under stress. The volcano plot in **Fig. 2b** presents differentially expressed genes from RNA-sequencing data: a total of 1,905 genes were differentially expressed (for an adjusted p-value < 0.05) out of the 11,982 genes with nonzero total read count. A heatmap of the 25 most up- and down-regulated genes with significant enrichment analysis categories annotated on the right-hand side is presented in **Fig. 2c**. ISR-activation for 24 h shows a trend for enrichment of genes related to ISR-activation (10/25 genes) and cell signaling (14/25 genes). Significant GO-terms for enrichment on downregulated genes after ISR-activation for 24h are related to the extracellular matrix (5/25 genes) as well as glial cell migration (3/25 genes).

**Figure 2.**
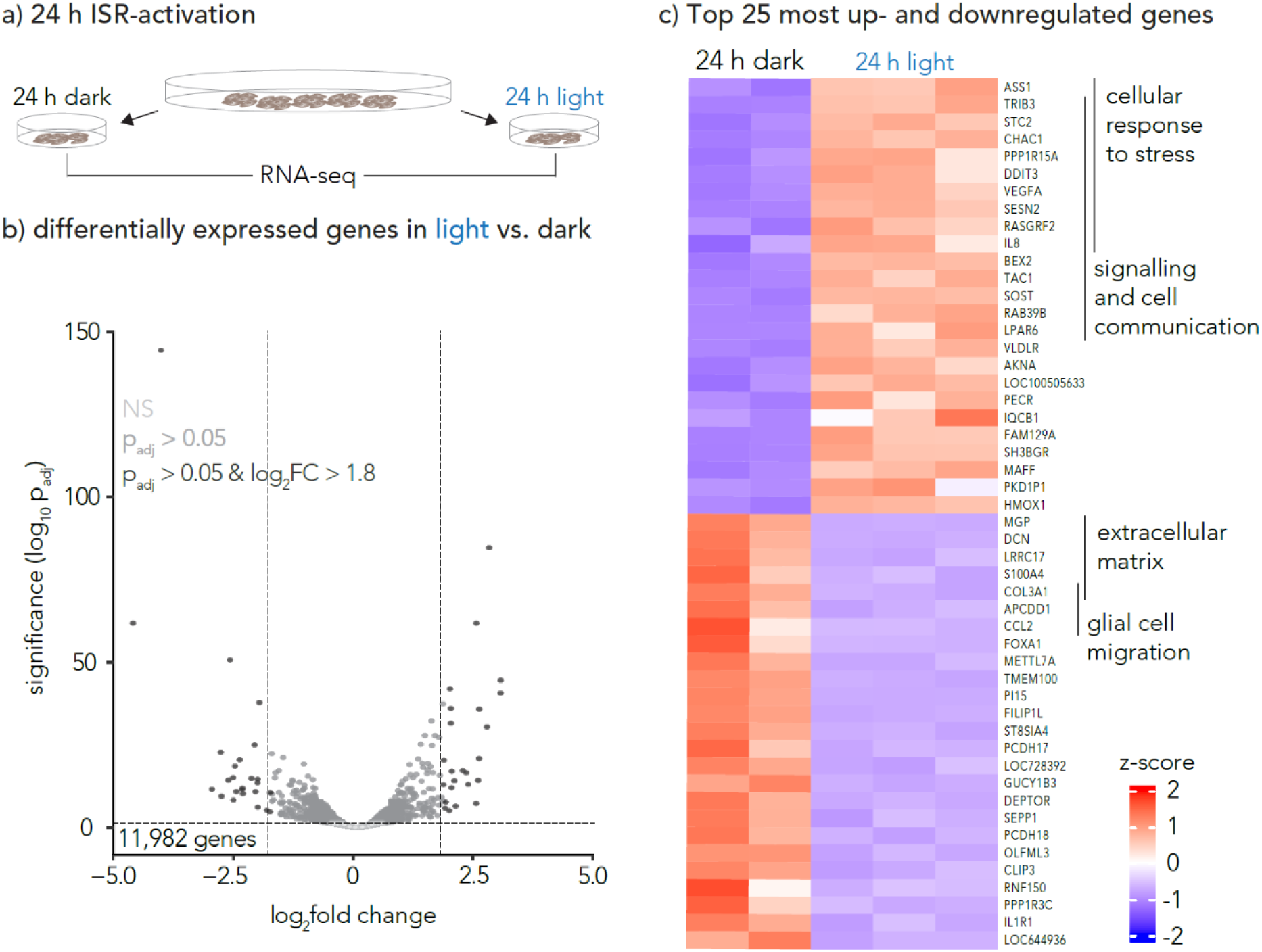
a) Differential gene expression analysis was performed after 24 h stimulation of opto-PKR H4 cells in light (N=3) compared to control cells 24 h in dark (N=2). b) Volcano plot presents differentially expressed genes from RNA-sequencing data: a total of 1,905 genes were differentially expressed out of the 11,982 genes with nonzero total read count, for an adjusted p-value < 0.05. Nonsignificant differentially expressed genes are colored in light gray, all differentially expressed genes with an adjusted p-value larger than 0.05 are colored dark grey, and genes with p-value criteria as well as a log_2_fold change larger than 1.8 are colored black. c) Heatmap of the 25 most up- and down-regulated genes with significant enrichment analysis categories annotated on the right-hand side. ISR-activation for 24 h shows a trend for enrichment of genes related to ISR-activation and cell signaling. Significant GO-terms for enrichment on downregulated genes after ISR-activation for 24 h are related to the extracellular matrix as well as glial cell migration.

In addition, single genes in the differentially expressed set were examined and some of the most pronounced were found to have tumor suppressor-nature, including for example, upregulation of *BEX2*, a reputed tumor suppressor gene commonly silenced in gliomas including GBM [104], [105], and downregulation of the *MGP* gene, encoding for protein crucial for cell migration [106]. Thus, at the gene-level, regulation of the extracellular space as well as the cancer cells’ migratory behavior indicate cancer regulation by the ISR.

### 3.3. ISR-activation Changes the Extracellular Environment

Enrichment analysis post differential gene expressions showing a significant downregulation of genes in the extracellular space indicate a decrease in the ISR-activated cells’ contribution to the extracellular environment. To further examine this result at the protein level a fluorescence assay was performed using Collagen Stain Kit staining for fibrillar collagen and non-collagenous ECM proteins, on cells that were cultured for the same amount of time but stressed for different periods (**Fig. 3a**). ISRIB, a small molecule inhibiting the ISR, was used in control experiments as well as wild type cells without light-inducible ISR-activation.

Primarily, we noticed that ISR-activation of cells over 72 h decreased cell viability, shown in **Fig. 3b** where the number of cells in the well was quantified by the area fraction of nuclei coverage in confocal images. The area fraction for opto-PKR cells (N=2 well repeats) in light (“opto light”) was visibly lower than controls: WT cells in light (“WT light”, N=2 well repeats), opto-PKR cells in dark (“opto dark”, N=2 well repeats), and opto-PKR cells in light with ISR-inhibitor ISRIB (“opto light +ISRIB”, N=2 well repeats).

**Figure 3.**
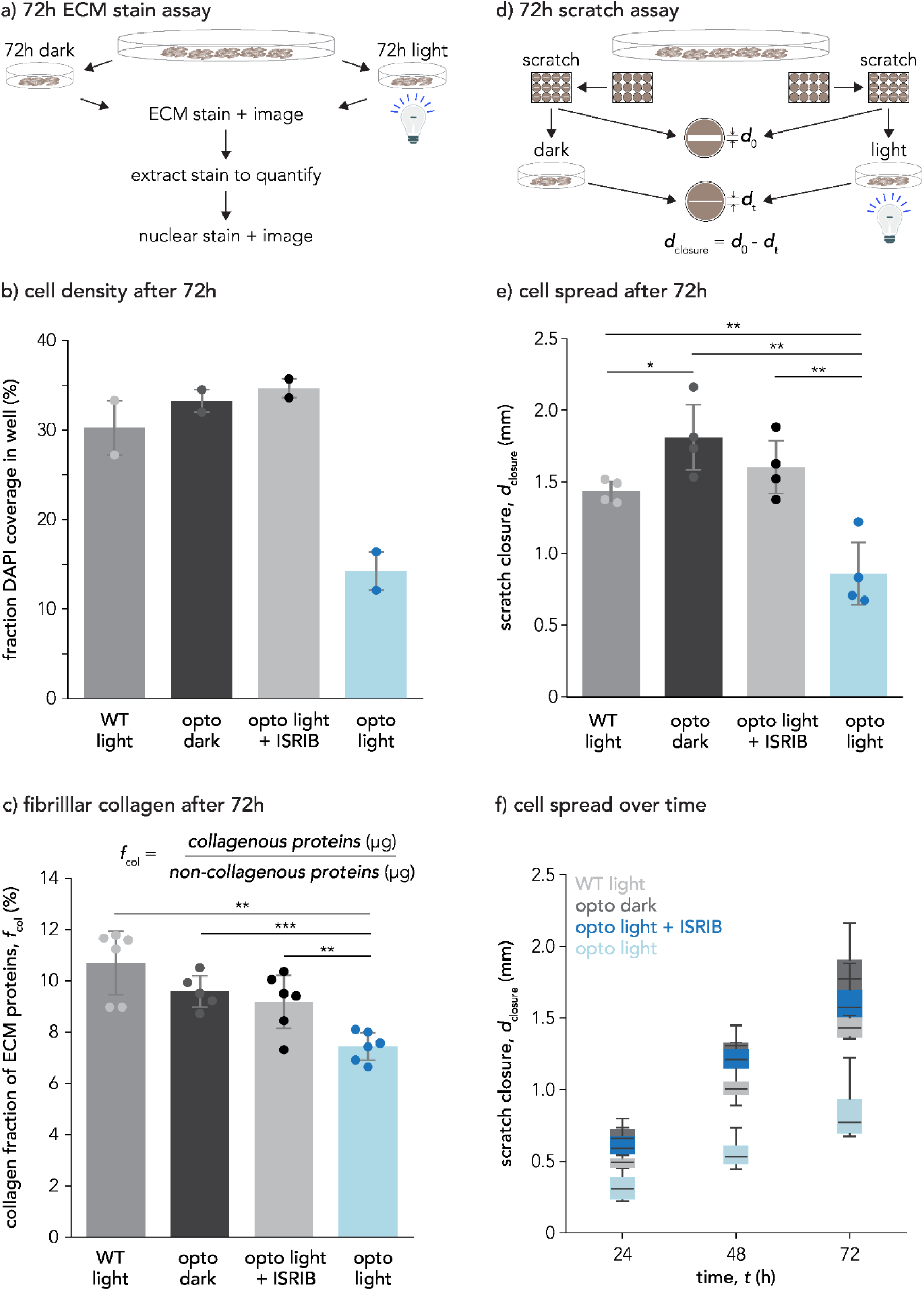
ISR-activation over 72 h shows reduced collagen content in the extracellular environment and inhibited cellular spread. In all sub-figures, opto-PKR H4 cells with ISR-activation (“opto light”) are compared to wild type cells in light (“WT light”), opto-PKR cells without ISR-activation (“opto dark”), and opto-PKR cells in light with inhibited ISR-activation (“opto light +ISRIB”). a) Procedure for 72 h ECM stain assay. b) Quantification of cell viability after 72 h of ISR-activation based on DAPI stain area coverage. Each data point represents quantification of a whole well in a 24-well plate. c) Quantification of fibrillar collagen to non-collagen ECM ratio after 72 h of stimulation. Each data point represents quantification of a whole well in a 24-well plate. d) Procedure for 72 h scratch assay. e) ISR-activation inhibits the spread of cells, quantified from scratch closure distance in scratch assay over 72 h (representative images shown in **Supplementary** Fig. 5). Each data point represents the average scratch closure in one well of a 12-well plate. f) Time-resolved quantification of cellular spread in scratch assay from 0 h, to 24 h, to 48 h, up to 72 h of stimulation compared to controls. For all the subfigures, boxplots and barplots indicate mean values with error bars showing standard deviations (* p < 0.05, ** p < 0.01, *** p < 0.001).

To account for the loss of cells over 72 h in the ECM stain assay, the amount of collagenous protein detected was normalized by the amount of non-collagenous protein detected for each condition. The fraction of collagen in the ECM is significantly reduced under ISR-activation (opto light, N=6 well repeats) compared to WT light (Mann-Whitney, p = 0.0022, N=6 well repeats), opto dark (t-test, p = 0.0003, N=5 well repeats), opto light +ISRIB (t-test, p = 0.0072, N=6 well repeats), supporting results from the differential gene expression analysis and gene enrichment analyses (**Fig. 3c**). The observed rescue of both cell viability as well as the amount collagenous proteins in the control condition where ISRIB is added to opto-PKR cells in light further supports that both cell death and collagen reduction is originating specifically from ISR-activation.

Representative images of wells stained with DAPI, Sirius Red, and Fast Green are shown in **Supplementary** Fig. 5.

### 3.4. ISR-activation Inhibits Spread of Cells in 2D

To further assess if ISR-activation does counteract cell migration as suggested by differential gene expression analysis, a scratch assay was performed (**Fig. 3d**). The results show significant inhibition of cellular spread with ISR turned on (opto light, N=4), compared to control conditions: WT light (t-test, p = 0.0047, N=4), opto dark (t-test, p = 0.0019, N=4), opto light +ISRIB (t-test, p = 0.0041, N=4), supporting results from the differential gene expression analysis and gene enrichment analyses (**Fig. 3e**). Additionally, we observed that the cellular spread also was reduced with longer time under stress from 24 h, to 48 h, to 72 h (**Fig. 3f**), again comparing opto light (N=4), to WT light (N=4), opto dark (N=4), and opto light +ISRIB (N=4). The observed rescue of cellular spread in the control condition where ISRIB is added to opto-PKR cells in light further supports that inhibited cellular spread is an effect of ISR-activation. Representative images from scratch assays are shown in **Supplementary** Fig. 5.

### 3.5. ISR-activation Promotes Containment of GBM Cells in 3D

Reduced spread in 2D with ISR-activation inspires control of GBM tumor containment in 3D. Expanding our 2D model into 3D culture of spheroids we aimed to study if these pathological alterations translate into a model that more closely resembles a tumor.

Spheroids were generated from WT and opto-PKR cells of the GBM cell line SF-295 in low-attachment 96-wells, thereafter, put into Collagen I matrix and imaged every hour over a day on a confocal microscope, while stimulated every 2 minutes with light or kept in the dark. ISR-activated spheroids (opto light) showed visibly increased containment compared to control conditions of WT cells in light (WT light) and opto-PKR cells in the dark (opto dark) (**Fig. 4a**). Within less than 12 hours, clear protrusions were formed in the control wells where spheroids were imaged in red light to not induce stress-activation (**Fig. 4b**). Quantification of maximum intensity projection images of z-stacks taken at every hour of the time lapse show steady increase of spheroid area change for the control conditions, while kept almost constant in the “opto light” ISR-activated spheroid (**Fig. 4c**). Another quantification of cellular spread in 3D is the spheroid perimeter change (**Fig. 4d**), where the ”opto light” condition has much less increase in perimeter change compared to the control conditions. However, the “opto light” perimeter change does increase slightly over time because of few small protrusions that still occur under stimulation. Additionally, spheroid circularity, *c*, was calculated. For a perfectly circular object *c*=1, meanwhile protrusions from spheroids drastically decrease *c* to a value less than one. For ISR-activated spheroids the circularity remained higher for longer time compared to control conditions (**Fig. 4e**). The few protrusions occurring in the “opto light” condition drastically changes the circularity *c*, as it scales inversely with the square of the perimeter *p*, why the circularity is not maintained at *c*=1 throughout the time lapse.

**Figure 4.**
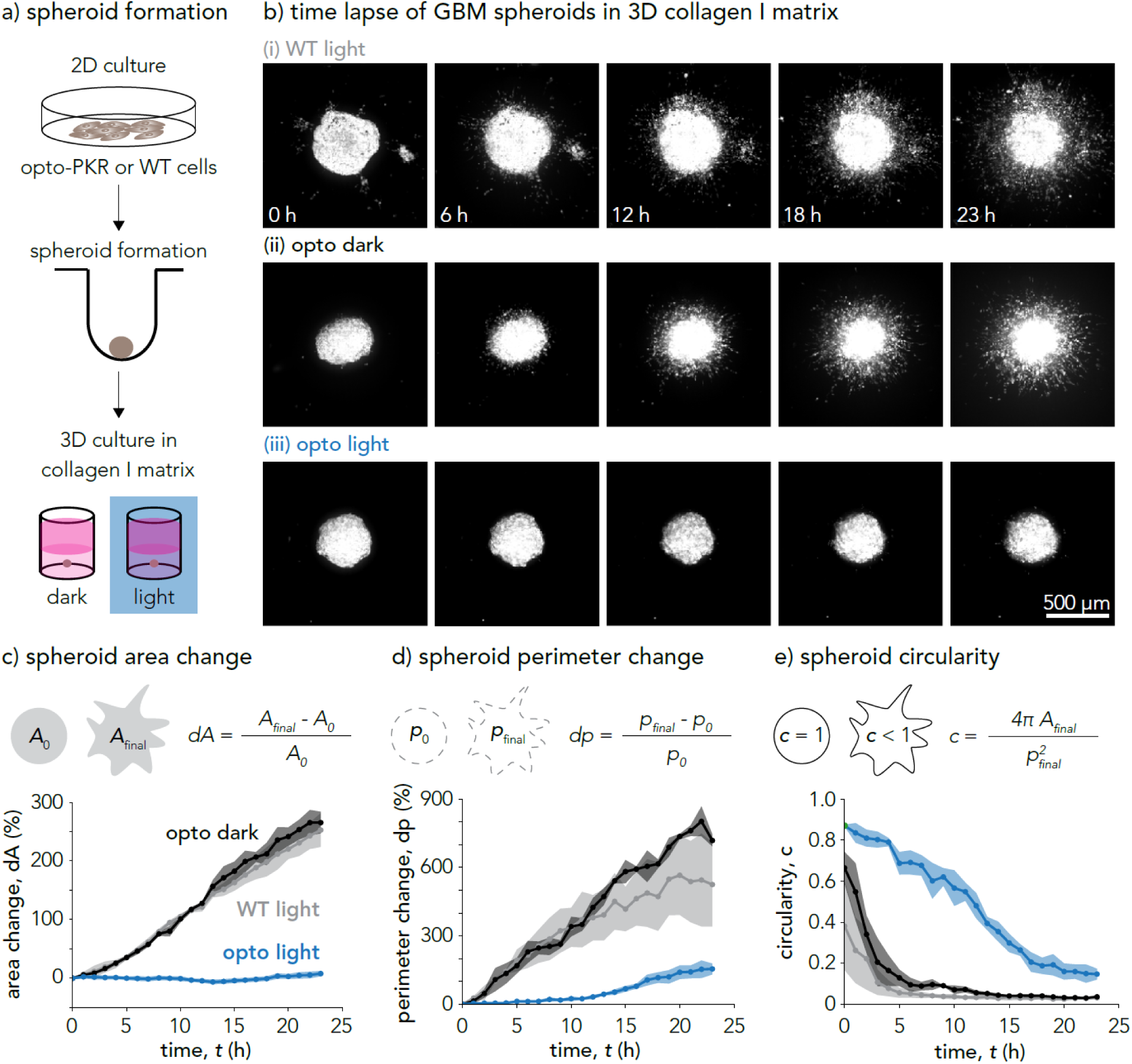
Light-controlled ISR-activation limits GBM spread in 3D. a) Spheroid formation protocol for 3D culture in collagen I matrix. b) Maximum intensity projection images of confocal z-stacks of GBM spheroids. Time lapse imaging over one day of ISR-activation in opto-PKR cells (opto light) compared to controls; opto-PKR cells in dark (opto dark), and WT cells in light (WT light). c), d), and e) Quantifications of spheroid containment by area change, perimeter change, and circularity. Circles show data points and shaded regions are standard deviations based on n=3 technical replicates. Images and corresponding quantifications in this figure are representative data from experiments on the cell line SF-295 dyed with CellTracker Red.

For non-time lapse experiments, spheroids were generated from six different cell lines (WT and opto-PKR cell lines for each of the three cell lines H4, SF-295, and U251) in low-attachment 96-wells, thereafter, put into Collagen I matrix and studied at time point 0 and after 24 hours on a confocal microscope. Significantly more and longer protrusions were formed in all control condition spheroids compared to the ISR-activated opto light spheroids (**Fig. 5a**). Quantification of maximum intensity projection images of z-stacks taken after 24 h compared to time point 0 shows steady increase of spheroid area as well as perimeter for all control conditions, compared the opto light ISR-activated spheroid (**Fig. 5b-d**). P-values for all conditions can be found in **Supplementary Table 1**.

**Figure 5.**
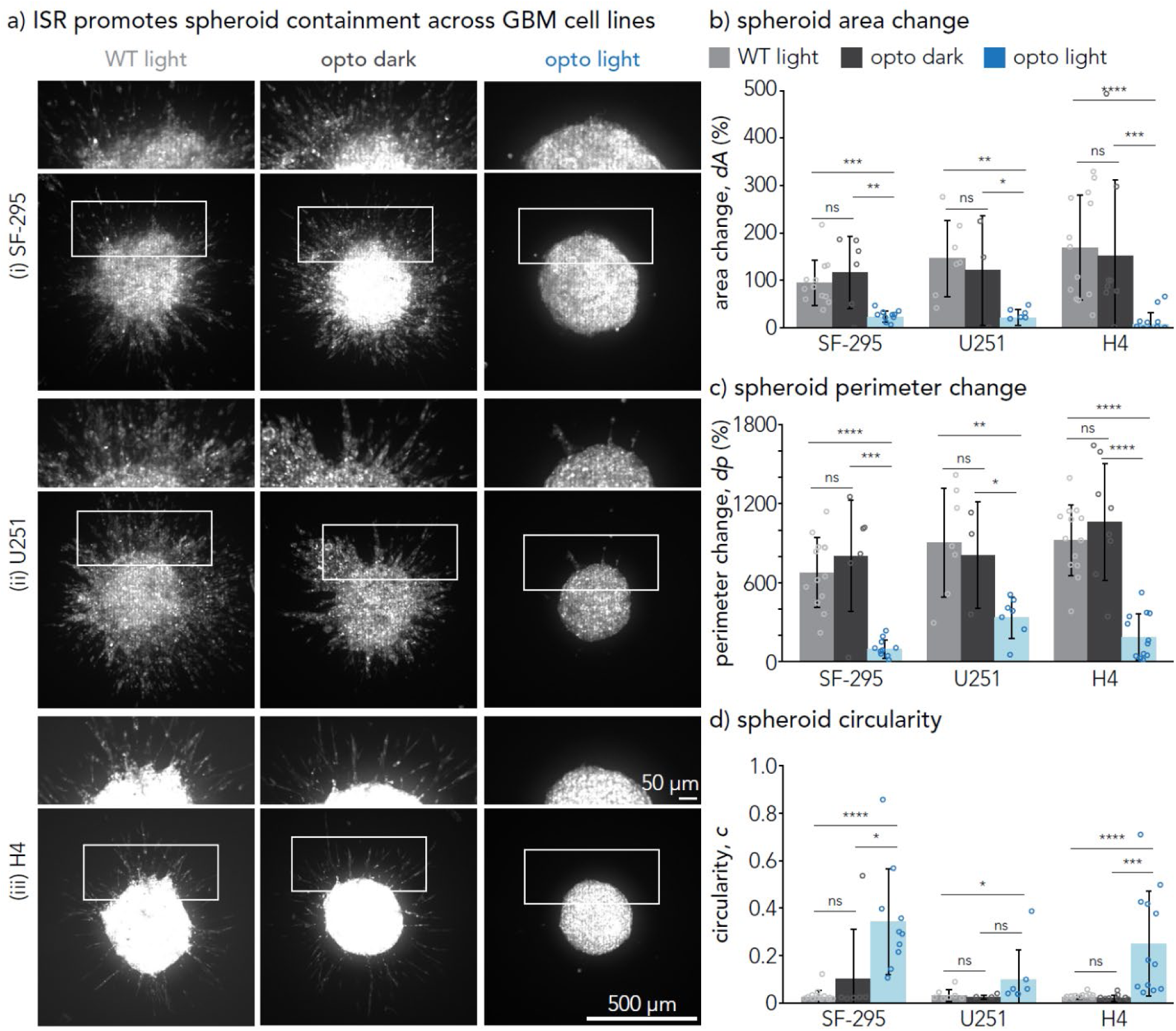
ISR-activation limits GBM spread in 3D across different cell lines (SF-295 GBM, U251 GBM, H4 neuroglioma). Spheroids in collagen matrix were imaged before and after stimulation with light or kept in the dark for 24 h. a) Maximum intensity projections from confocal imaging of spheroid z-stacks after 24 h in collagen matrix. Controls (WT light, opto dark) are compared to ISR-activated spheroids (opto light). Figure b), c) and d) show quantifications of spheroid containment after 24 h from maximum intensity projections of z-stacks from confocal imaging. Boxplots show mean values, with error bars indicating standard deviations. Each data point represents one spheroid. All cell lines examined show significant containment under ISR-activation compared to controls (* p < 0.05, ** p < 0.01, *** p < 0.001, **** p < 0.0001).

Additionally, we observed some differences in protrusion morphology between the cell lines (e.g., H4 neuroglioma protrusions tend to be less dense than SF-295 GBM protrusions), as can be seen in **Fig. 5a**. Altogether, these results suggest that ISR-activation can regulate GBM containment towards less aggressive tumor spread.

### 3.6. Photopatterned GBM Containment with Precise Spatiotemporal Control of ISR-activation

Reduced GBM spread in 3D with ISR-activation poses the question whether the migratory effect is intrinsic or extrinsic to the cancer cells. To investigate this, we employed photopatterned ISR-activation using digital micromirror devices to focus blue LEDs onto part of the spheroid.

Spheroids were generated from opto-PKR cells of the GBM cell line SF-295 in low-attachment 96-wells, thereafter, put into collagen I matrix and imaged every hour over a day on a confocal microscope, while only one half of the spheroid was stimulated every 2 minutes with light or kept in the dark. The ISR-activated side of the spheroid (light, right hand side) showed visibly more containment compared to the side of the spheroid kept in the dark (dark, left hand side) (**Fig. 6a**). Quantification of maximum intensity projection images of z-stacks taken at every hour of the time lapse show steady increase of spheroid area change for the side of the spheroid kept in the dark, while kept almost constant in the illuminated, ISR-activated part of the spheroid (**Fig. 6b**). These results indicate that spread inhibition in the GBM cells may be an intrinsic response from the cells rather than communicated and signaled to surrounding cells, based on the observations at the length- and time scales studied.

**Figure 6.**
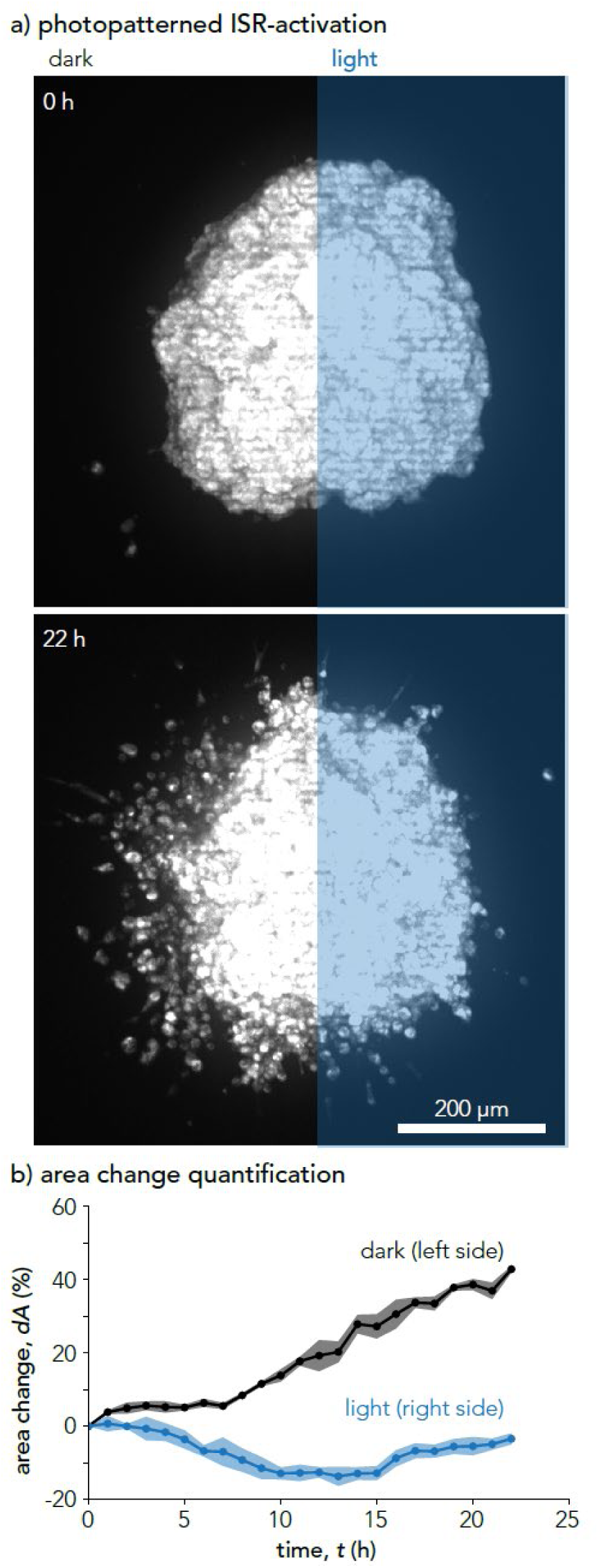
Photopatterned GBM containment in SF-295 spheroid obtained with precise spatiotemporal control of ISR-activation. a) Using digital micromirror devices on a confocal microscope, only a rectangular region of interest was illuminated to spatially constrain induction of ISR-activation to one half of a spheroid. b) Quantification of spheroid containment by the area change for each of the two half-spheres pictured in a) (light vs. dark) over 22 h. Circles show data points and shaded regions are standard deviations based on n=3 technical replicates. The spheroid was dyed with CellTracker Red.

### 3.7. Spheroids Recover Spreading Post ISR-activation

As a cell is relieved from stress by an external stressor or pathological condition, recovery may be expected depending on the intensity and duration of the stress experienced. To examine recovery in our system, we activated the ISR for 24 h, and thereafter kept them in the dark to recover, resulting in protrusion formations forming already within the next 48 h across all three cell lines (**Fig. 7**).

**Figure 7.**
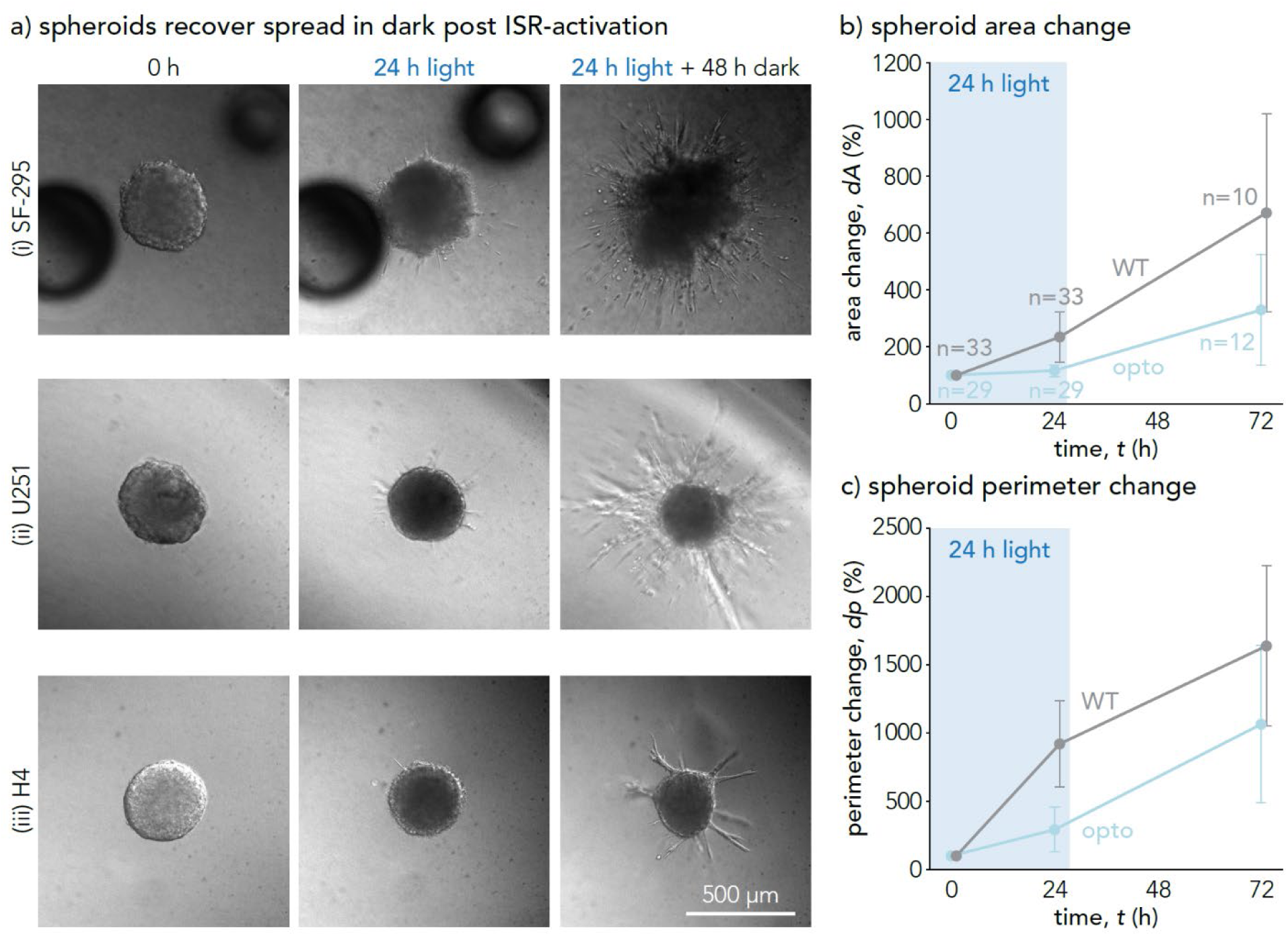
Spheroids recover spread after ISR-activation across multiple cell lines. To study recovery after ISR-activation, opto-PKR spheroids were illuminated for 24 h, imaged, thereafter kept in the dark for 48 h and imaged again. a) DIC images of opto-PKR spheroids at timepoints 0 h, 24 h and 72 h. b) and c) Quantifications of spheroid spread recovery across all cell lines (SF-295, U251, H4) WT cells compared to opto-PKR cells.

## 4. Discussion

The role of ATF4 in GBM progression has in multiple recent studies seemed to be a promotor of GBM progression [78], [79], [81], [82], [85], [86], while others have shown GBM suppression, including ISR-activation from the drug Nootkatone [84], Withaferin A [83], and specific ISR-activators used together with temozolomide [87], [88], [89]. Among the studies showing GBM suppression with ISR-activation, they all had in common the upregulation of CHOP together with ATF4. Thus, we seek to clarify the role of ISR-activation with both ATF4 and CHOP upregulation in GBM cells.

This study is the first to examine the ISR’s role in GBM with optogenetics, avoiding possible side effects from drugs. The results presented in this work, including a range of different experimental modalities (e.g., IF time courses, RNA-sequencing, ECM-stains, 2D scratch assay, as well as 3D spheroids) all trending the same direction suggesting that light-induced ISR-activation via opto-PKR, including activation of both ATF4 and CHOP, changes the extracellular environment as well as suppresses aggressive cancer spread. Our results indicate that this effect might be cell intrinsic based on spatial control at the tissue scale. Furthermore, the recovery study indicate that ISR-activation might support a tunable, non-ablative intervention space, which would be beneficial for translational applications. These findings suggest a route to containment and motivate ISR-activating small molecule screening in GBM models.

There is a variety of known ISR-inducing small molecules (e.g., Salubrinal, Halofungione, Sephin1, and recently discovered ISR-potentiators [107], [108], [109], [110]) that could be screened *in vivo* models for GBM cancer treatment. Especially interesting might be to study such drugs as candidates for delivery together with common GBM treatments [89], like the work of Weatherbee *et al.* where the drug JLK1486 was used for ISR-activation together with temozolomide [88].

Reversely, Jain *et al.* took advantage of the aggressive migratory behavior in GBM and developed nanofibers successfully guiding the tumor cells from an inoperable site to an extracortical location [111]. As ISR-activation inhibits GBM migration, such tumor-guiding materials may benefit from an ISR-inhibitor like ISRIB to minimize the number of cells with potential migration inhibition due to stress.

Following recovery post ISR-activation, a fascinating aspect of the ISR yet to be well understood, is so called stress memory, potentially an underlying mechanism for stress resilience and rapid organ and cell adaptations [112]. Stress memory can be studied as the change in response to a second incident of stress based on previous stress exposure, exemplified by *C. elegans* studies where, for example, oxidative stress at young age improved the organisms’ resistance to heat shock later in life [113], [114]. Stress memory has also been simulated and predicted in theoretical models, where the interplay between positive and negative feedback, especially GADD34, in the ISR seem to be a key regulator [90], [115].

Cellular structure is another factor suggested to play an important role in the ISR, as actin dynamics have been shown closely linked to its negative feedback system. When eIF2α is phosphorylated in the activation of the ISR, the transcription factor GADD34 is sequentially elevated for negative regulation of the ISR [5], [6]. The GADD34 will together with PP1 and, more recently proven, G-actin form a complex that de-phosphorylates the eIF2 [7], [8]. The actin cytoskeleton in the cell is a continuously dynamic component where globular actin, or G-actin, can polymerize into filamentous actin, F-actin, and the process is reversible [116]. As more G-actin polymerize into F-actin, there are less monomers available for de-phosphorylation of eIF2 to regulate an activated stress response, which has been shown using the drug jasplakinolide, known to stabilize F-actin polymers [7], [117]. The effect of ISR-activation has been found different between different studies across multiple cancers and cell types, and the reason behind remains unclear [118]. Further studies would be needed to investigate differences in ISR-activation between cell types and potential relations to their structure.

As actin dynamics relates to both extracellular cues and the control mechanisms of the ISR, the extracellular environment may play a role in tuning ISR-dynamics and stress memory. Studies of cellular spread and migration on different culture platforms have shown differences in the actin structure between cells on soft and stiff substrates and have also been seen to differ between different cell types [119]. Thus, stiffness of different cell culture substrates may influence ISR-dynamics and stress memory.

As stress induces downregulation of genes related to the extracellular environment, what does that tell us about the preferred tumor microenvironment for containment? To better understand what the role the extracellular space has in GBM containment, identification of crucial components for migration recovery studies of the ECM in relation to the cellular spread would help understand the time scales and relevancy. Could the extracellular environment function as a “memory structure” for a cell after ISR-activation, and possibly influence recovery times post stress?

In this work, collagen I was used as a model system for a 3D tumor microenvironment for GBM. While the brain microenvironment does not contain large amounts of collagen I, previous work has shown that this environment has been one of the more triggering for creating long finger-like protrusions with single cells migrating long distances of GBM cells [120], [121], [122], [123]. We herein show that the ISR can inhibit aggressive spread in a GBM-protrusion triggering 3D model environment.

While efforts were put forward to quantify the containment of GBM spheroids, the precision of doing so to obtain absolute values remains a challenge. Various segmentation algorithms were examined for the job, while in the end manual outlining of the spheroids gave the most accurate quantification across samples because of the highly complex morphology of the structures. This method is not perfect, which can be seen from particularly in perimeter change standard deviations from multiple technical replicates, but for the interest of relative comparison between images rather than absolute values the method used shows quantifications mirroring the visual difference between the images taken.

## 5. Concluding Remarks

In this work, an optogenetic tool was used to examine the influence of the ISR on GBM. Our results suggest that ISR-activation to a level including ATF4 and CHOP-activation changes the extracellular environment as well as suppresses aggressive spread of this cancer. We show that this effect might be cell intrinsic based on spatially defined regions of less cellular spread in photopatterned ISR-activation at tissue scale. We also observe time dependent effects, where recovery of finger-like protrusions is seen within days after ISR-activation. These findings open a new venue for potential therapeutic targets in GBM treatment, including ISR-activating compounds. In addition, an optogenetic tool like the one used herein is well suited to further study the effect of ISR-activation across different types of cancer, as well as other diseases, to better understand the growing role this complex signaling network seems to play in many of today’s most challenging diagnoses.

## Acknowledgements

This work was primarily supported by the ICB (Award No. W911NF-19-2-0026). AAP acknowledges funding support from the NSF CAREER award (CMMI-CAREER-2048043), from which LKM was supported by over summer 2025. MZW and ED were supported by the US Army Research Office under contract W911NF-19-D-0001 and cooperative agreement W911NF-19-2-0026 for the Institute for Collaborative Biotechnologies. LH was supported by the California Institute for Regenerative Medicine under the COMPASS Training Award (EDUC5-13744).

