## Supplementary Information for "Optogenetic Control of the Integrated Stress Response Limits Glioblastoma Invasion"

[1]Materials Department
[2]Interdisciplinary Program in Quantitative Biosciences
[3]College of Creative Studies, Biology
[4]Molecular, Cellular, and Developmental Biology
[5]Center for BioEngineering
[6]Neuroscience Research Institute
University of California Santa Barbara, Santa Barbara, California, USA 93106

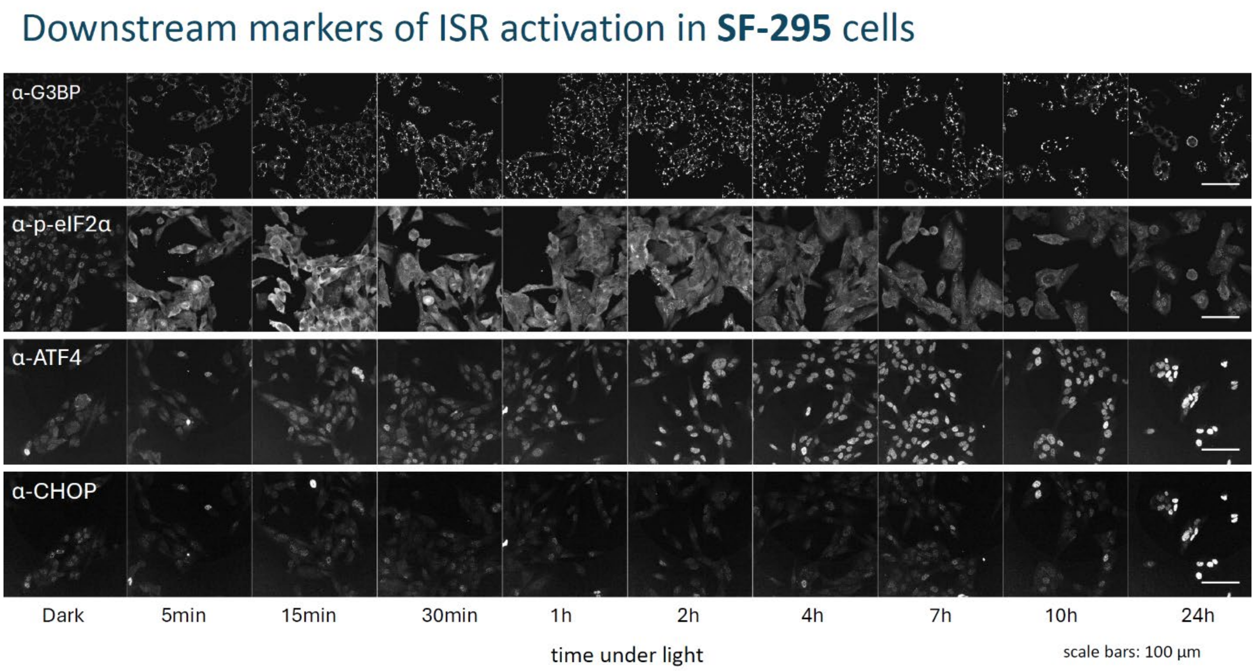


**Figure 1** Immunofluorescence imaging of fixed and stained SF-295 opto-PKR GBM cells in 24h light (pulsed 5s ON, 15s OFF) compared to time zero for proteins involved in ISR-activation: phosphorylation of eIF2a, G3BP (in stress granules), ATF4, and CHOP. Quantification of immunofluorescence is shown in **Fig. 1** in the main manuscript.

### Downstream markers of ISR activation in **U251** cells

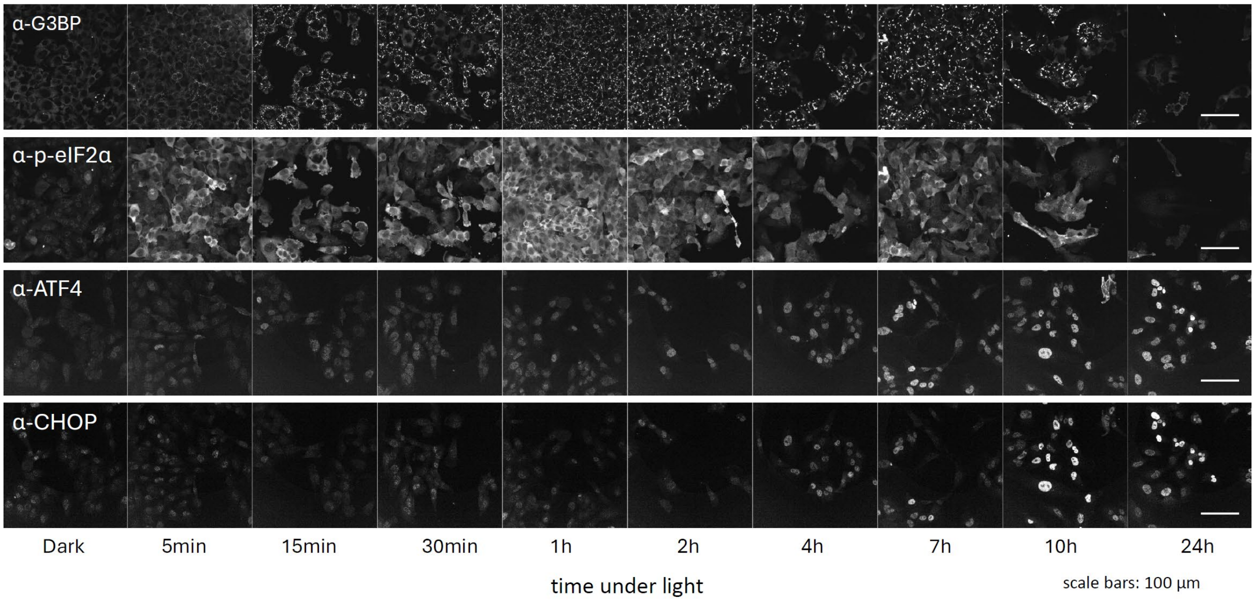


**Figure 2** Immunofluorescence imaging of fixed and stained U251 opto-PKR GBM cells in 24h light (pulsed 5s ON, 15s OFF) compared to time zero for proteins involved in ISR-activation: phosphorylation of eIF2a, G3BP (in stress granules), ATF4, and CHOP. Quantification of immunofluorescence is shown in **Fig. 1** in the main manuscript.

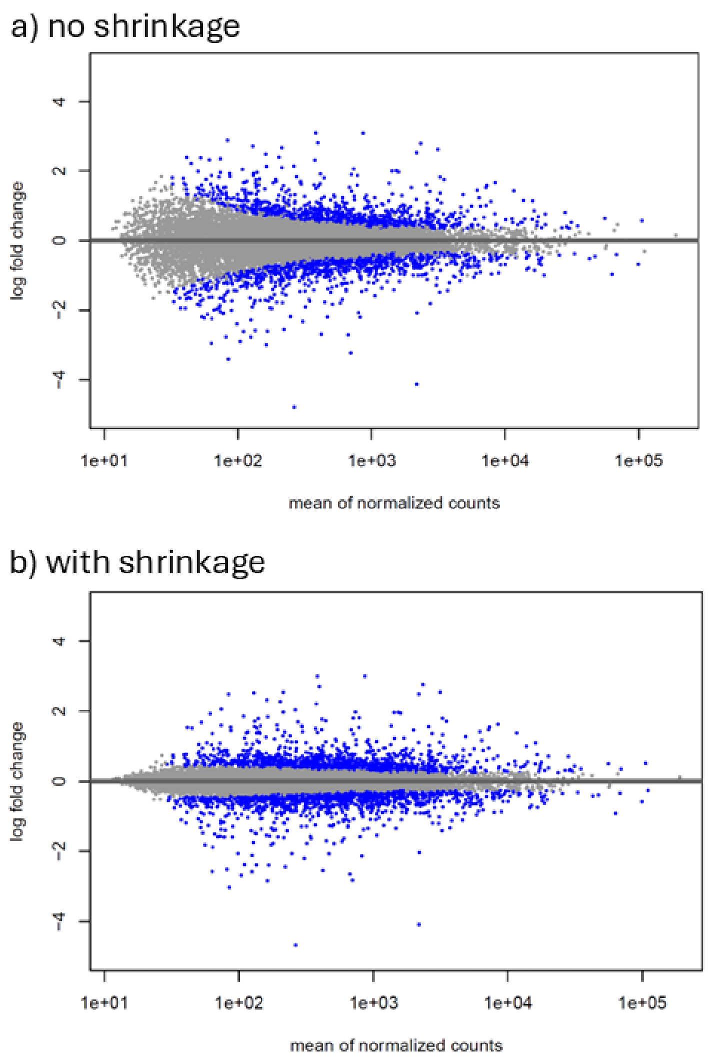


**Figure 3** MA plots for all differentially expressed genes from RNA-sequencing of opto-PKR H4 cells in 24h light compared to 24h dark-samples before (a) versus after (b) shrinkage using *LFCshrink* (correction for high log fold change values on low count samples to avoid false positive) [1].

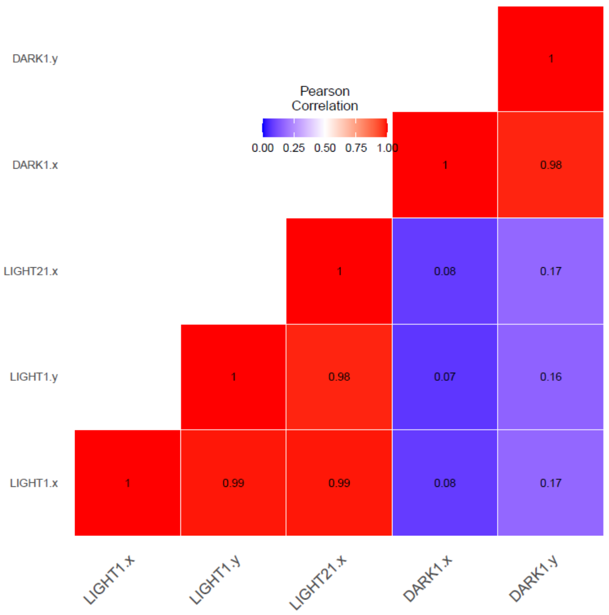


**Figure 4** Pearson correlation heatmap between RNA-sequencing samples.

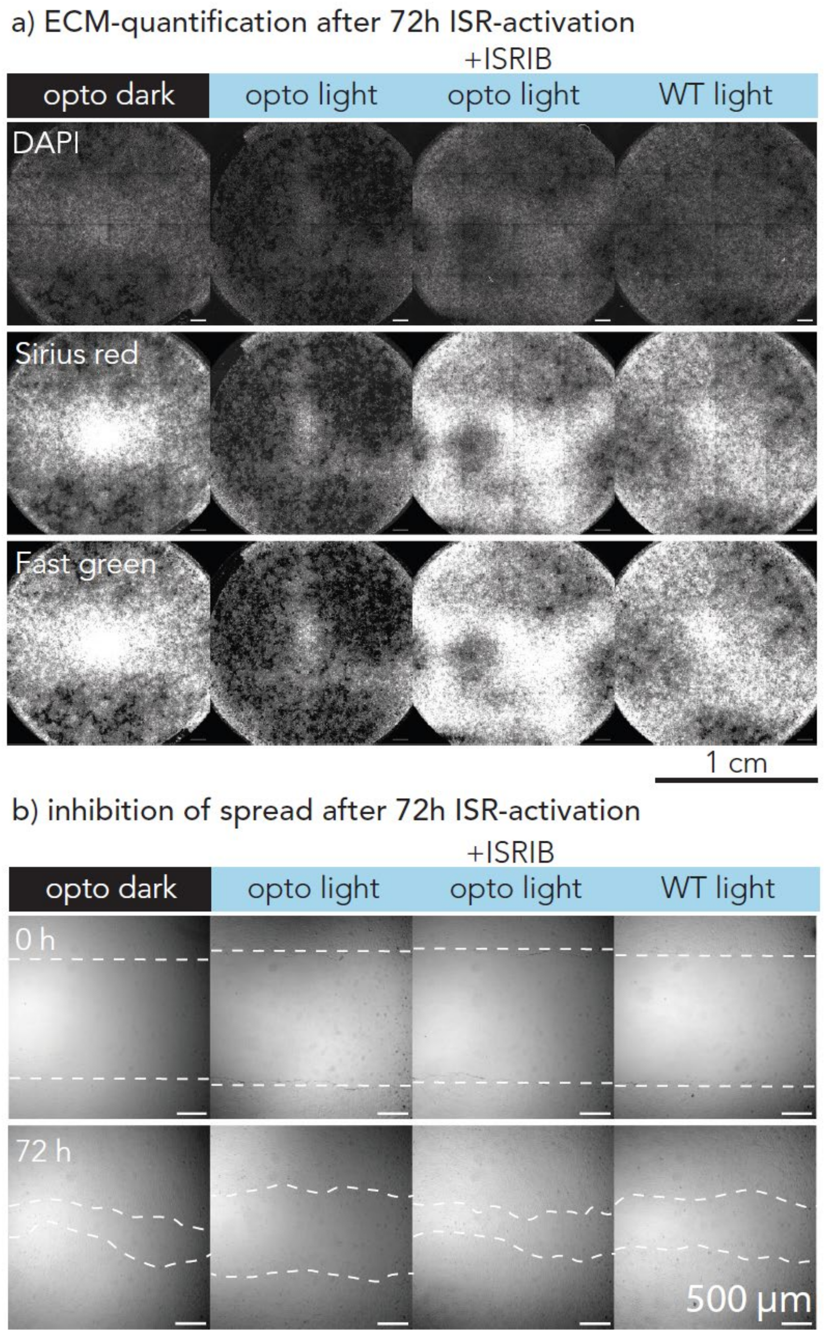


**Figure 5** a) Representative images of whole wells with H4 cells with nuclear stain (DAPI), as well as ECM stain (Sirius Red - collagenous protein, Fast Green - non-collagenous protein). b) Representative images of scratch assay for quantification plots in main **Fig. 3**.

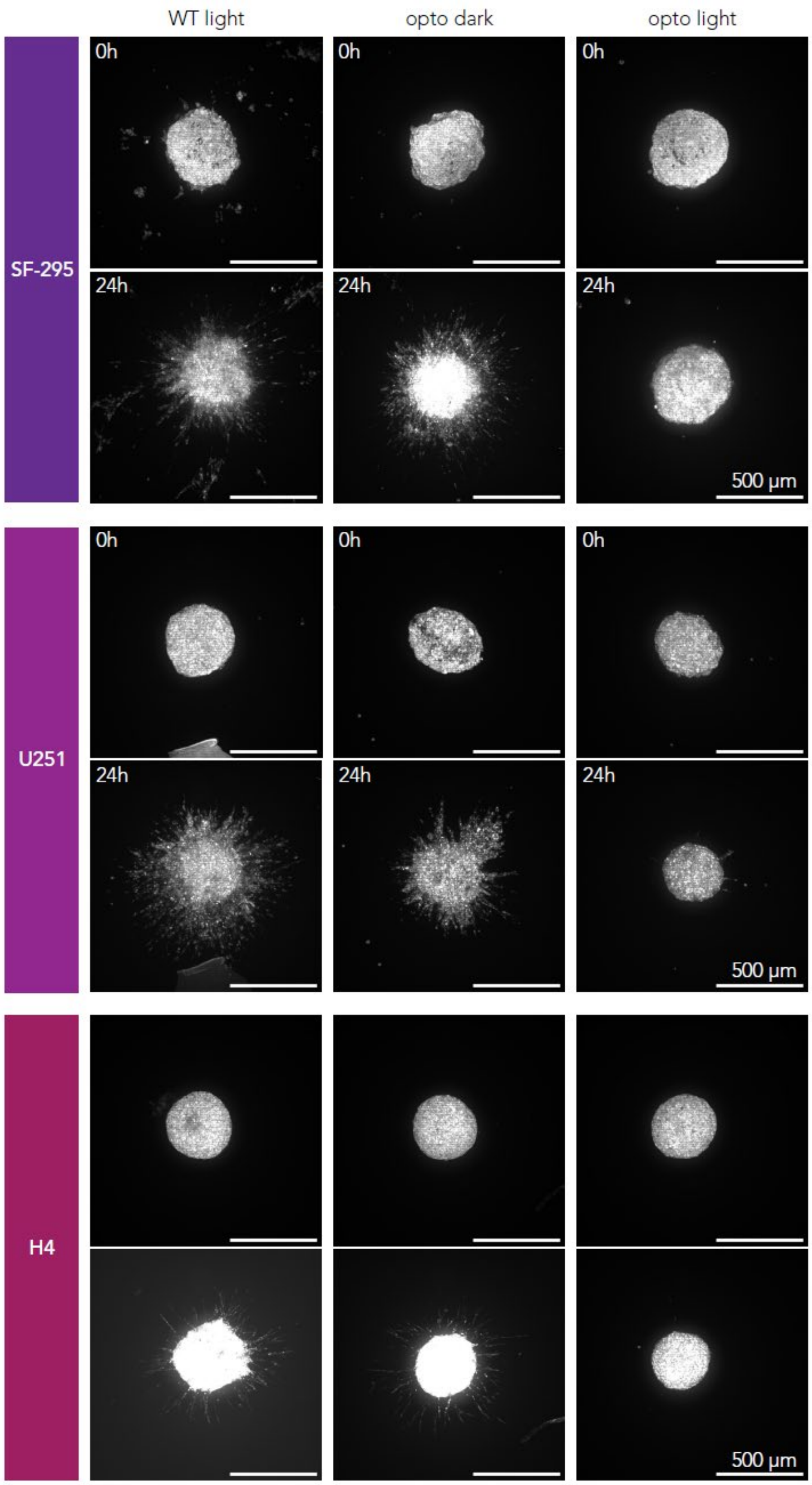


**Figure 6** Representative images for each cell line (SF-295, U251, H4) showing spheroid containment after one day (24h) of ISR-activation compared to controls. These images are the same as in **Fig. 5** in the main manuscript but include images for each spheroid at time point 0h.

| Cell line | Parameter | Comparison | Shapiro test | Signif. test | p-value | # spheroids |
|---|---|---|---|---|---|---|
| SF-295 | Area change | opto light/WT light | 0.85/0.11 | t | 1.4e-4 | N=10/N=13 |
| SF-295 | Area change | opto light/opto dark | 0.85/0.19 | t | 1.8e-3 | N=10/N=6 |
| SF-295 | Area change | WT light/opto dark | 0.11/0.19 | t | 0.45 | N=13/N=6 |
| U251 | Area change | opto light/WT light | 0.69/0.92 | t | 1.9e-3 | N=7/N=7 |
| U251 | Area change | opto light/opto dark | 0.69/0.64 | t | 0.043 | N=7/N=3 |
| U251 | Area change | WT light/opto dark | 0.92/0.64 | t | 0.70 | N=7/N=3 |
| H4 | Area change | opto light/WT light | 0.015/0.10 | MW | 6.4e-5 | N=12/N=13 |
| H4 | Area change | opto light/opto dark | 0.015/0.012 | MW | 7.1e-4 | N=12/N=8 |
| H4 | Area change | WT light/opto dark | 0.10/0.012 | MW | 0.75 | N=13/N=8 |
| SF-295 | Perimeter change | opto light/WT light | 0.69/0.91 | t | 1.3e-6 | N=10/N=13 |
| SF-295 | Perimeter change | opto light/opto dark | 0.69/0.21 | t | 1.1e-4 | N=10/N=6 |
| SF-295 | Perimeter change | WT light/opto dark | 0.91/0.21 | t | 0.43 | N=13/N=6 |
| U251 | Perimeter change | opto light/WT light | 0.43/0.78 | t | 5.2e-3 | N=7/N=7 |
| U251 | Perimeter change | opto light/opto dark | 0.43/0.38 | t | 0.022 | N=7/N=3 |
| U251 | Perimeter change | WT light/opto dark | 0.78/0.38 | t | 0.76 | N=7/N=3 |
| H4 | Perimeter change | opto light/WT light | 0.10/0.84 | t | 4e-8 | N=12/N=13 |
| H4 | Perimeter change | opto light/opto dark | 0.10/0.85 | t | 7.5e-6 | N=12/N=8 |
| H4 | Perimeter change | WT light/opto dark | 0.84/0.85 | t | 0.38 | N=13/N=8 |
| SF-295 | Circularity | opto light/WT light | 0.17/3.12e-5 | MW | 8.1e-5 | N=10/N=13 |
| SF-295 | Circularity | opto light/opto dark | 0.17/2.9e-5 | MW | 0.020 | N=10/N=6 |
| SF-295 | Circularity | WT light/opto dark | 3.12e-5/2.9e-5 | MW | 0.48 | N=13/N=6 |
| U251 | Circularity | opto light/WT light | 2.0e-4/8.1e-3 | MW | 0.04 | N=7/N=7 |
| U251 | Circularity | opto light/opto dark | 2.0e-4/0.11 | MW | 0.085 | N=7/N=3 |
| U251 | Circularity | WT light/opto dark | 8.1e-3/0.11 | MW | 0.65 | N=7/N=3 |
| H4 | Circularity | opto light/WT light | 0.050/0.019 | MW | 3.9e-5 | N=12/N=13 |
| H4 | Circularity | opto light/opto dark | 0.050/0.011 | MW | 3.8e-4 | N=12/N=8 |
| H4 | Circularity | WT light/opto dark | 0.019/0.011 | MW | 0.10 | N=13/N=8 |

**Table 1** p-values from significance tests on quantifications of spheroid geometry after one day (24h) of ISR-activation vs. controls from **Fig. 5** in the main manuscript. Controls (WT light, opto dark) are compared to opto-PKR spheroids in light “opto light” (from cell lines SF-295, U251, H4) illuminated with light. Shapiro test was used to determine if data sets were normally distributed. MW = Mann-Whitney, t = t-test.
